# Attention Samples Items in Visual Working Memory Rhythmically

**DOI:** 10.1101/2022.04.12.488012

**Authors:** Samson Chota, Carlo Leto, Laura van Zantwijk, Stefan van der Stigchel

## Abstract

Attention allows us to selectively enhance the processing of specific locations or features in our external environment while simultaneously filtering out momentarily irrelevant information. It is currently hypothesized that this is achieved through the boosting of relevant sensory signals which biases the competition between competing neural representations. Recent neurophysiological and behavioral studies have revealed that attention is a fundamentally rhythmic process, tightly linked to neural oscillations in fronto-parietal networks. Instead of continuously highlighting a single object or location, attention rhythmically alternates between multiple relevant representations at a frequency of 3 – 6 Hz. However attention can not only be directed towards the external world but also towards internal visual working memory (VWM) representations, e.g. when selecting one of several search templates to find corresponding objects in the external world. Two recent studies have revealed that objects in VWM are attended in a similarly rhythmic fashion as perceived objects. We add to the current literature by showing that retro-cues towards multi-feature gratings in VWM initiate a similar theta-rhythmic competition, modulating reaction times in an anti-phasic manner. Our findings add to the converging body of evidence that external and internal visual representations are accessed by highly similar, rhythmic attentional mechanisms.

## Introduction

While our subjective visual experience might seem like a continuous stream of incoming information, accumulating evidence suggests that perception as well as attention undergo rhythmic fluctuations. When attending two locations in our visual field, reaction times to unpredictable targets are modulated as a function of time at a theta frequency (Fiebelkorn et al., 2013; Holcombe & Chen, 2013; Landau & Fries, 2012; Song et al., 2014), potentially as a result of neural competition (Chota et al., 2018; Kienitz et al., 2018). The periodic modulations of performance are observed for each location individually but are often found to oscillate in antiphase, congruent with a rhythmic spotlight of attention that samples multiple locations alternatingly and sequentially (Fiebelkorn & Kastner, 2019; VanRullen, 2013). Not only does this seem to be the case for spatial locations but also for non-spatial features such as color or orientation (i.e., feature-based attention (Mo et al., 2019; Re et al., 2019a).

Attention directed towards stimuli or features in the external world is assumed to facilitate processing at least partially via boosting of sensory signals (Boynton, 2009; Brefczynski & DeYoe, 1999; Corbetta et al., 1990; Gandhi et al., 1999; Martínez et al., 1999; Reynolds & Heeger, 2009; Somers et al., 1999). A similar mechanism might underly the behavioral benefits observed when directing attention towards internal (memorized) representations in visual working memory (VWM), commonly described as the retro-cue effect. Predictive cues directed towards one or more items in working memory have been shown to increase recall performance, even if cues are presented several seconds after encoding (for a review see Souza & Oberauer, 2016). A growing number of studies have revealed a strong overlap in the neurophysiological correlates of attention towards external (currently perceived) stimuli and internal working memory processes (Gazzaley & Nobre, 2012). This strong overlap is further supported by findings that strongly implicate early sensory cortices in the maintenance of working memory representations often referred to as the sensory recruitment hypothesis (Christophel et al., 2017; Ester et al., 2016; Gayet et al., 2018; Iamshchinina et al., 2021; Lorenc et al., 2021; Rademaker et al., 2019; Scimeca et al., 2018). For instance, attentional selection of items in working memory has been shown to bias activity in sensory areas that process these stimuli (Capilla et al., 2014; Poch et al., 2014) and to modulate decodability of item features most strongly in early visual cortices (Christophel et al., 2018).

To summarize, both external visual stimuli and internal VWM representations are encoded at least partially by overlapping neural populations on which attention needs to act. This raises the intriguing possibility that similar or even identical attentional mechanisms might be engaged for external and internal visual representations. Recent work has begun to extend the rhythmic nature of internal attention directed towards VWM items. Peters et al. instructed participants to memorize four dots that defined the endpoints and hence the outlines of 2 objects (Peters et al., 2018). A subsequent retro-cue directed and reset attention to one of these objects. At densely-sampled (200 ms to 1000 ms) but unpredictable timepoints a target was presented that fell within or outside of the cued or un-cued object. Peters et al. observed that reaction times (RT) measured as a function of the distance to the cue were rhythmically fluctuating at a frequency of 6 Hz. Furthermore, the RT time-series of responses to the cued versus the un-cued object were modulated in antiphase, indicating that attention periodically sampled the two object locations in VWM sequentially. A second study conducted by Pomper et al. showed that similar fluctuations in performance occurs between two oriented bars in VWM (Pomper & Ansorge, 2021). Importantly, they presented their stimuli sequentially and at fixation, reducing potential influences of rhythmic spatial attention. The efficiency of probe processing fluctuated at a frequency of 6 Hz and an anti-phasic relationship between the accuracy time-series of the first and second stimulus was observed. Interestingly, they found that the appearance of the second stimulus did not always reset the attentional rhythm acting on the first item, potentially due to a lack of an explicit external cue.

The recent emergence of studies finding rhythmic behavioral fluctuations in internal attention, closely resembling those found in external attention, suggest a close relationship between both attentional mechanisms and a central involvement of rhythmic brain activity. A potential mechanistic overlap could have wide-ranging implications for our understanding of how the brain selects internal and external information and how this selection is orchestrated by oscillatory activity. Importantly rhythmic attentional fluctuations in working memory have only been shown for items consisting of single features. However attention has been strongly implicated in supporting binding of multiple features both during perception and in working memory (Hitch et al., 2020; Treisman, 1996). Here we set out to further characterize the behavioral correlates of theta-rhythmic attentional sampling in working memory using items consisting of multiple relevant features in a retro-cue paradigm. We hypothesize a rhythmic modulation of the efficacy of memory-probe comparisons at a frequency of 3 to 6 Hz. Furthermore, we hypothesize that efficacy time-series of both memory items will be modulated in anti-phase, indicative of an alternating attentional sampling of working memory representations.

## Materials and Methods

### Participants

26 participants (aged 19-28, 10 females) with normal or corrected to normal vision participated in the experiment. Informed consent forms were signed before the experiment. After completion participants were compensated with academic credits or 12 euro. The experiment was carried out in accordance with the protocol approved by the local ethical committee and followed the Code of Ethics of the World Medical Association (Declaration of Helsinki).

### Experimental Design

Stimuli were presented at a distance of 58 cm with a LCD display (27 inch, 2560 × 1440 resolution, 120 Hz refresh rate) using the Psychophysics Toolbox^i^ running in MATLAB (MathWorks). Stimuli consisted of a central fixation cross (diameter = 0.3°), two circular memory stimuli consisting of randomly oriented gratings (diameter = 10°; spatial frequency = 4 cpdva) both of which had a random color selected from opposite phases of the CIELAB color space (48 sampled colors; L = 59, a = 18, b = - 8), a central color cue (0.93°), a probe (diameter = 10°; spatial frequency = 4 cpdva) whose orientation was shifted slightly relative to the same colored memory stimulus, and a mondrian mask consisting of a colored pattern (diameter = 10°; code from (Christophel et al., 2012)).

Participants performed a 2FA working memory task in which they were instructed to compare the orientation of two memory stimuli to the orientation of a probe that matched the color of one of them. Behavior in response to the probe was measured using a dense-sampling approach that allowed us to measure the rhythmic effect of attentional fluctuations on the processing of the probe. Trials began with the presentation of the fixation cross on a blank screen for 1000 ms (Figure 1). This was followed by the sequential presentation of two memory stimuli for 1000 ms each (Stimulus A and Stimulus B). After a blank delay of 1000 ms we presented a cue for 250 ms that matched the color of either Stimulus A or Stimulus B and indicated which of the two working memory items will be probed with 75% validity. Based on previous studies we hypothesized that the cue would reset rhythmic attentional sampling while still requiring participants to maintain a representation of the un-cued item (Fiebelkorn et al., 2013; Landau & Fries, 2012; Mo et al., 2019; Peters et al., 2018). The cue was followed by a blank screen of variable delay (between 200 ms and 700 ms in steps of 16.6 ms) after which we presented a Probe stimulus for 75 ms that matched either Stimulus A or Stimulus B in color and was oriented slightly clock-wise or counter clock-wise. After another brief delay of 25 ms we presented a mask for 75 ms after which participants responded if the orientation of the Probe was tilted clock-wise or counter clock-wise relative to the same-colored WM stimulus. Participants responded by pressing either the Q (counter clock-wise) or the P button (clock-wise).

**Figure 1.**
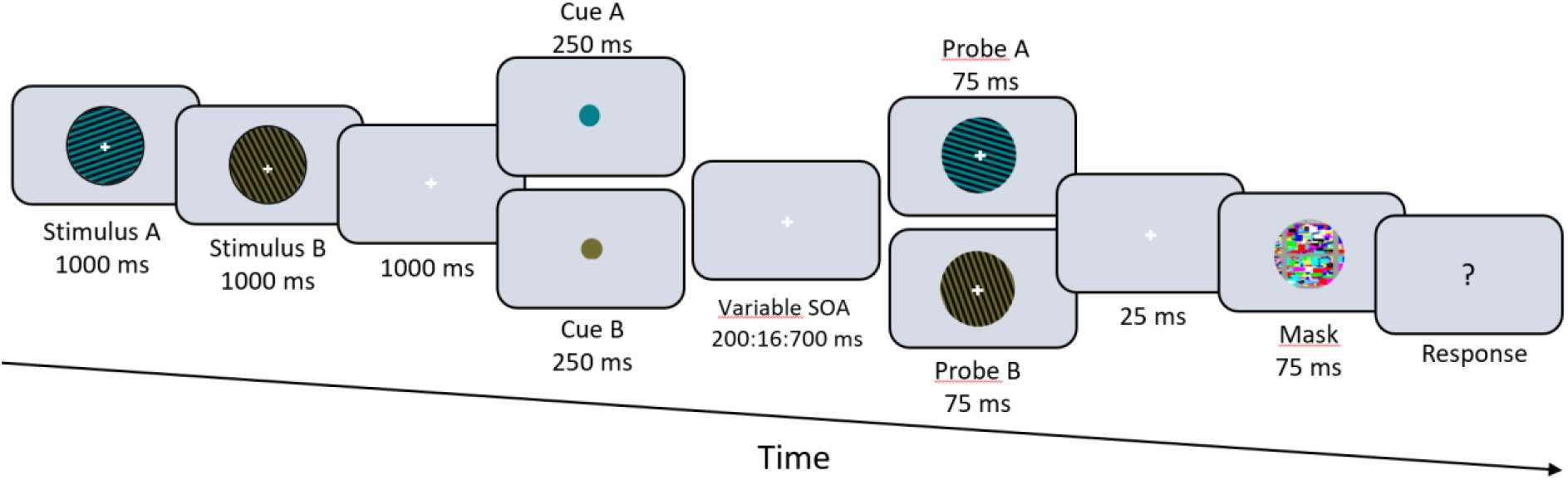
Task paradigm. Participants encoded two gratings with different colors and orientations. A subsequent retro-cue, indicative of the probe color (75% validity), was presented to reset attention to one of the items in working memory. We measured behavior at several densely sampled timepoints (between 200 and 700 ms) by presenting the probe and asking participants if it was rotated clock-wise or counter clock-wise relative to the same-colored item in working memory.

The experiment consisted of a familiarization block (∼4 minutes) in which participants performed several trials with binary feedback on every trial. This was followed by a staircase procedure (QUEST, 80 trials, ∼10 minutes) that allowed us to tailor the angular difference in orientation between memory item and probe so that performance was on average at 75% (Watson, Pelli, 1983). During this procedure participants received binary feedback at the end of every trial. After completion of the staircase procedure the main experiment began (20 blocks of 30 trials each, ∼75 minutes). The main experiment was identical to the staircase procedure with the only exception being that feedback was now provided at the end of every block in the form of percent correct instead of at the end of every trial. This was done to minimize training effects. During the main experiment we continued to use the QUEST algorithm to keep task difficulty at 75% by dynamically adapting the angular difference between sample and probes.

### Data Analysis

We only considered trials with reaction times shorter than 2 seconds. For two participants this led to the exclusion of more than 20% of the trials and they were hence excluded from the analysis. For another participant our RT criteria led to the removal of all trials in a specific SOA and they were also removed from the analysis. 23 participants were included in the final analysis. To investigate rhythmic fluctuations in behavior we averaged reaction times within 30 different SOA’s for each individual participant to compute RT time-series. Only correct trials were considered since they most likely reflect successful working memory maintenance. As we hypothesized that the cue would reset attentional sampling to the cued item, irrespective of its serial position, we calculated the RT time-series for validly cued probes and invalidly cued probes separately. Individual RT time-series were detrended by fitting and removing a 2^nd^ order polynomial and moving averages were calculated using a 66.7 ms window. We then z-scored the RT time-series and applied a hamming window before performing spectral analysis using fast fourier transformation. The resulting individual power-spectra were averaged and we statistically tested for oscillatory peaks using a non-parametric permutation test: We repeated the above described preprocessing procedure 10.000 times, however in every iteration we randomly shuffled the SOA-RT pairs for every participant. This procedure allowed us to estimate the distribution of power-spectra under the null hypothesis that no rhythmic fluctuation is present in individual RT time-series. The real power-spectrum was then compared to the surrogate distribution of power-spectra on a frequency by frequency basis. Frequencies at which the real power-spectrum exceeded the 95% percentile of the surrogate distribution were considered significant as this equates to an alpha level of 0.05% (Figure 2CD). We performed a similar analysis on the difference wave between valid and invalid RT time-series (Figure 2E). Since we were mainly interested in low frequency fluctuations we only considered frequencies between 1 and 15 Hz.

**Figure 2.**
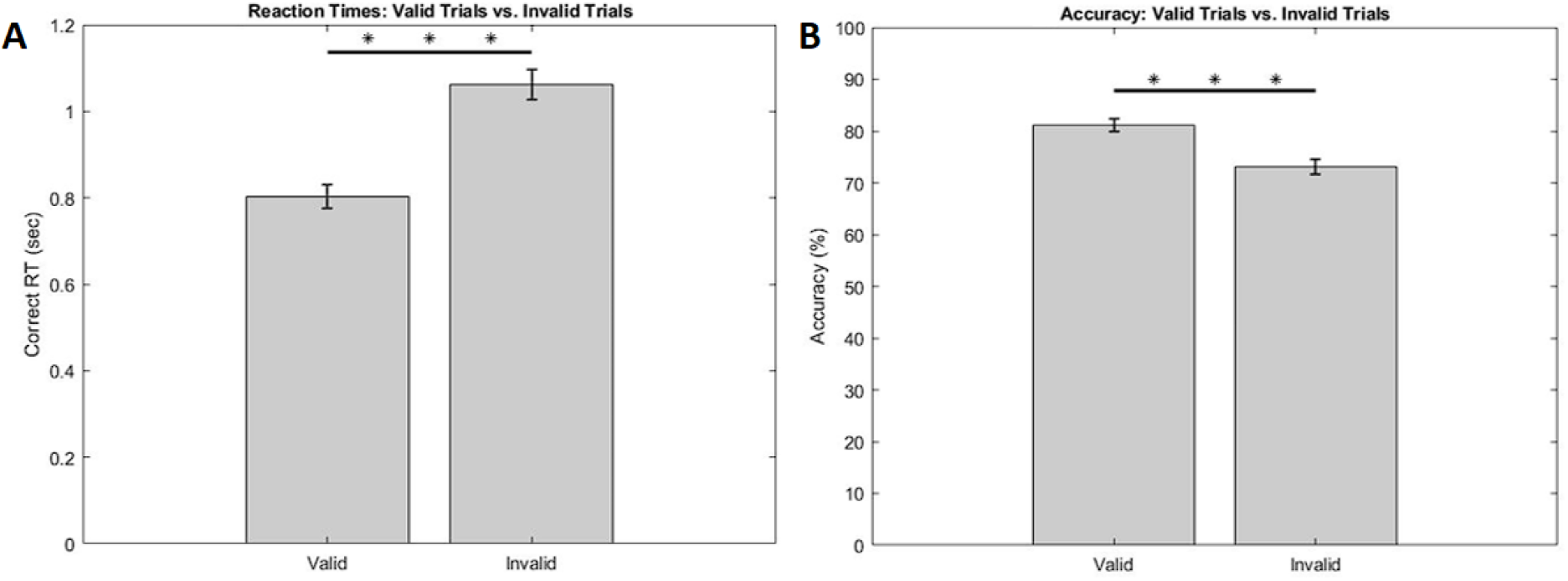
Average reaction times (B) and accuracy (A) for valid and invalid conditions

Oscillatory phases at 4.6 Hz were extracted from the complex components of the fourier transform of the individual preprocessed RT time-series. To investigate whether individual valid and invalid time-series were fluctuating in antiphase we subtracted the phase values and performed a circular t-test to compare average phase-difference values to 180° (Figure 2B).

Group average reaction times and accuracy were analysed using two sample t-tests.

## Results

Statistical analysis of the group average RT revealed that participants responded significantly faster to cued (mean = 831 ms) versus un-cued probes (mean = 1084 ms), t(22) = -12.97, p < .001). Furthermore, we found that reaction times were shorter when participants responded to probes matching the second stimulus (mean = 840 ms) as compared to the first stimulus (mean = 880 ms, t(22) = 3.85, p < .005). Similarly, accuracy was higher for cued probes (mean = 81.1 %) than for uncued probes (mean = 73.2%, t(22) = 8.76, p < .001) and higher for probes matching the second stimulus (mean = 81.4%) as compared to the first stimulus (mean = 76.8%, t(22) = 6.85, p < .001).

To investigate whether working memory representations are attended in a theta rhythmic fashion, we measured participants reaction times to matching probes at several densely sampled timepoints. We hypothesized that RT’s should be modulated at the theta rhythm, indicative of rhythmic fluctuations of the corresponding item in working memory. Furthermore, we hypothesized that RT time-series for responses to valid and invalidly cued probes should be modulated in antiphase, suggesting that attention periodically alternates between the two items in working memory.

To investigate rhythmic fluctuations in individuals responses we calculated RT time-series by averaging RT’s in each of 30 SOAs for responses to validly and invalidly cued items respectively. The spectral analysis of individual RT time-series revealed a significant peak at 3.75 and 4.6 Hz in the valid condition and no significant peaks in the invalid condition. Because of differences in the number of collected trials for valid (75%) and invalid (25%) conditions we might simply lack the statistical power to detect rhythmicities in the invalid condition. In addition, a recent similar study on behavioral oscillations has also failed to find a significant peak in the power-spectra of one of two conditions, revealing however that both conditions were still fluctuating in antiphase when analyzing individual oscillatory phases (Pomper & Ansorge, 2021). We therefore investigated if our two conditions were in antiphase by 1) Subtracting valid and invalid RT time-series, as the difference wave should reveal stronger oscillations (than each individual time-series alone) at a specific frequency when the two rhythms are in antiphase and 2) Analyzing phase difference between both conditions, as the difference should point towards 180 Hz if the oscillations are modulated in antiphase.

The spectral analysis of the valid minus invalid difference wave revealed a significant peak at 4.6 Hz and 5.6 Hz that was notably higher as compared to the valid condition (Figure 3C, peak at 0.025 db vs Figure 3E, peak at 0.06 db). Furthermore, our phase analysis of individual valid and invalid time-series revealed that the 4.6 Hz phase difference between valid and invalid trials was not significantly different from 180°, indicating an anti-phasic relationship of the oscillatory 4.6 Hz components (Figure 3B).

**Figure 3.**
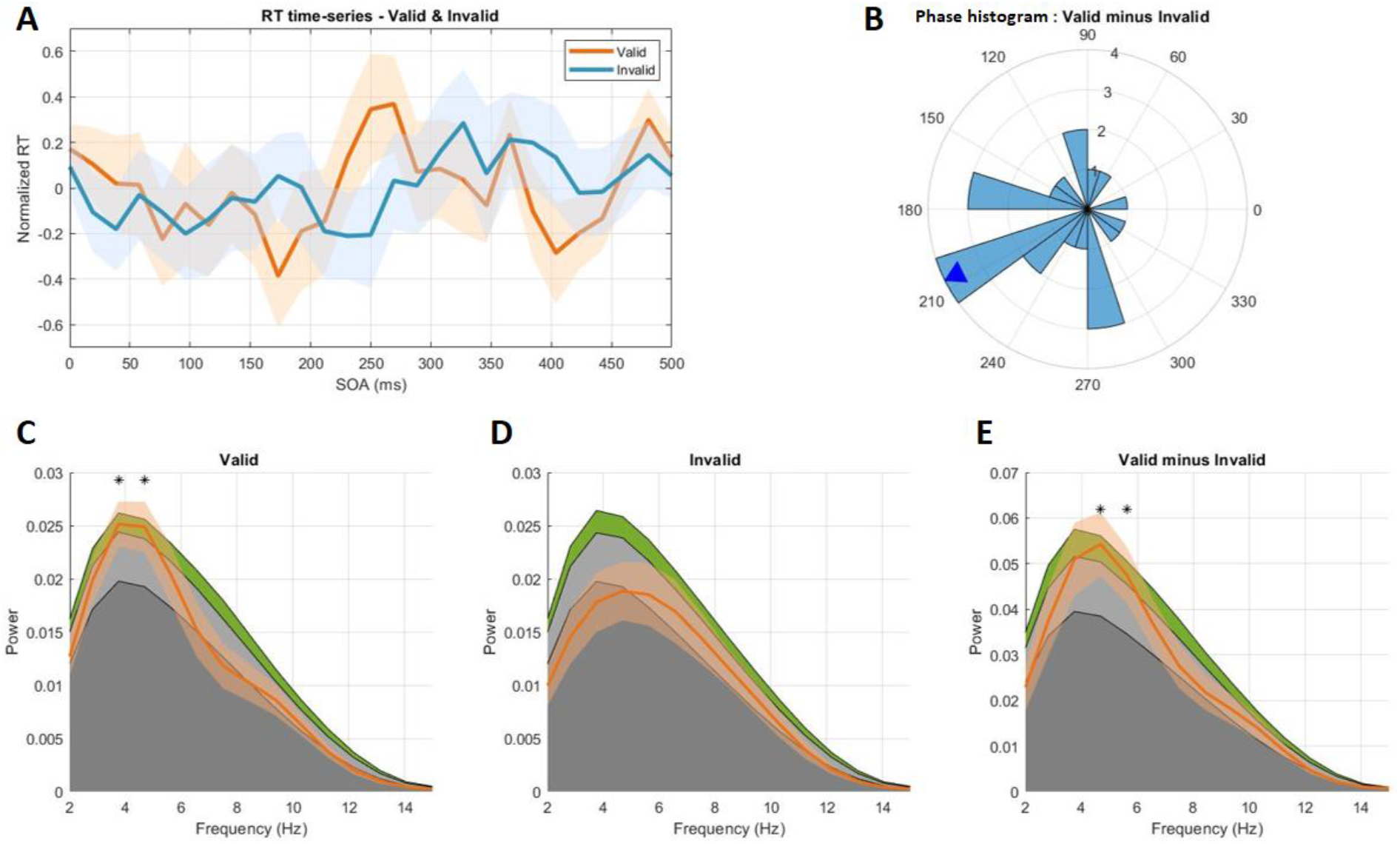
**A**. Group average RT time-series for valid and invalid trials. **B**. 4.6 Hz Phase difference plot. **C**. Power-spectrum RT time-series valid condition. Colored areas in the background display quantiles of statistical null distribution (dark grey: Power<Mean of permutation distribution, light grey: Power < 95% confidence interval, green: Power < 99% confidence interval). Stars indicate frequencies at which observed spectrum exceeded 95% of samples from permutation distribution. **D**. same as C but RT time-series of Invalid condition. **E**. Same as C and E but Valid minus Invalid RT time-series.

**Figure 4.**
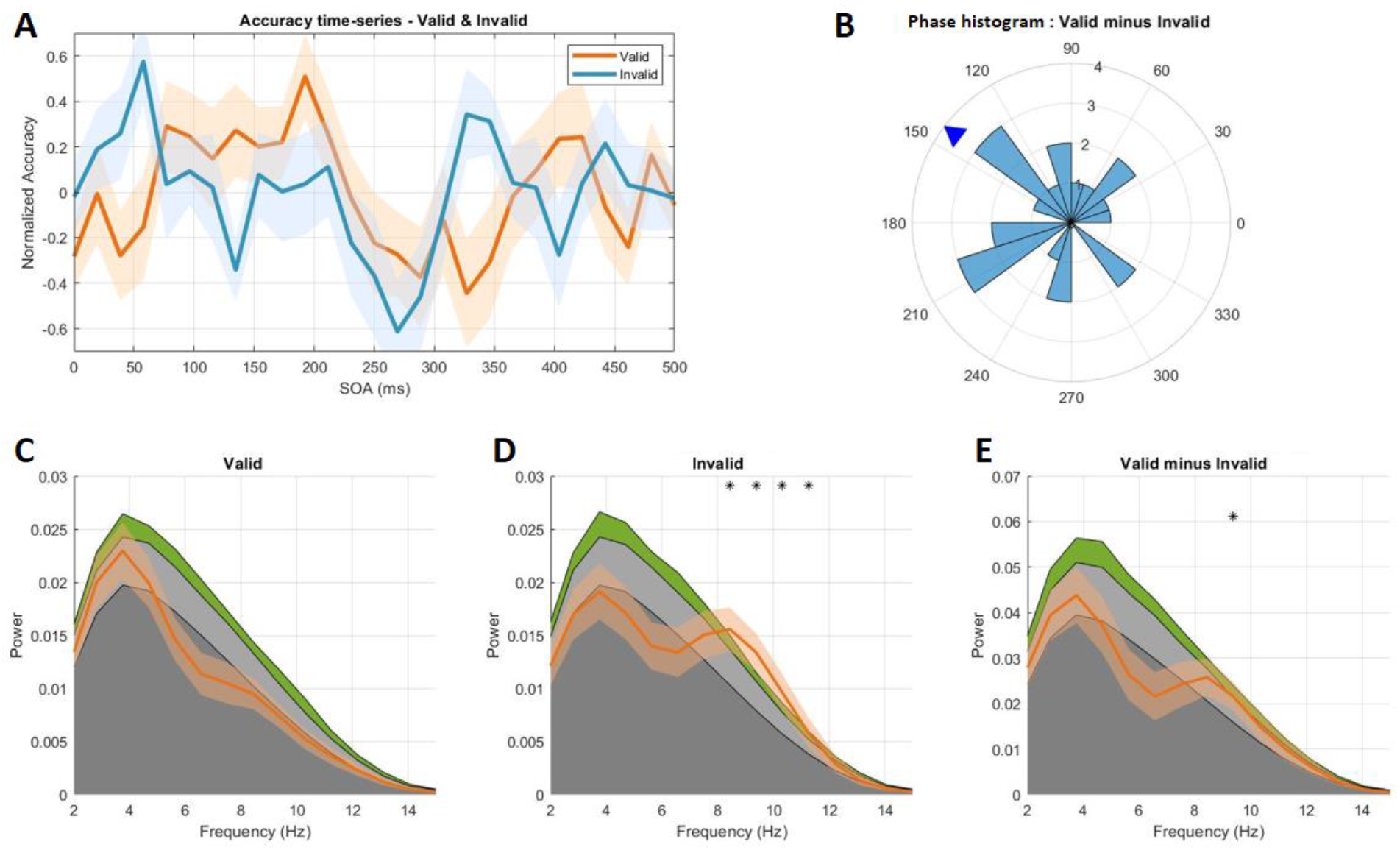
**A**. Group average accuracy time-series for valid and invalid trials. **B**. 4.6 Hz Phase difference plot. **C**. Power-spectrum accuracy time-series valid condition. Colored areas (dark grey: mean of permutation distribution, light grey < 95% confidence interval, green <99% confidence interval). **D**. same as C but accuracy time-series of Invalid condition. Stars indicate frequencies at which observed spectrum exceeded 95% of samples from permutation distribution. **E**. Same as C and E but Valid minus Invalid RT time-series.

We investigated rhythmic fluctuation in accuracy time-series in an identical fashion by comparing observed average power-spectra with stochastic null-distributions. The spectral analysis of individual accuracy time-series revealed a significant peak at 8.4, 9.4 and 10.3 Hz in the invalid condition. Although we found a significant peak at 9.4 Hz in the difference between valid and invalid accuracy time-series, the phase difference between both conditions was significantly different from 180° indicating no anti-phasic relationship. We subsequently applied Raley’s test of non-uniformity which revealed that phase differences were randomly distributed around the unit circle (p = 0.34) and hence had no consistent temporal relationship.

## Discussion

Spatial as well as feature-based attention to visually available information has been demonstrated to undergo rhythmic fluctuations. Here we set out to test if attention to features in working memory is modulated in a similar theta rhythmic manner. We find that a) the speed of responses to cued probes that match one of two items in working memory is modulated at a frequency of ∼4 Hz and b) rhythmic fluctuations at ∼4 Hz to cued and un-cued items are modulated in antiphase. Our results are indicative of a rhythmic and alternating attentional sampling of both working memory representations, which in turn modulates response speed to subsequent probes.

### Theta-rhythmic fluctuations in reaction times

While rhythmic fluctuations in RT time-series were observed for valid trials, we did not find a significant oscillatory peak in trials where participants were invalidly cued. However, when analyzing the individual 4.6 Hz phase of valid and invalid RT time-series we found a significant anti-phasic relationship. This suggests that an oscillatory component was present in invalid trials which our statistical analysis failed to reveal. We propose several possible reasons for this. First, since cue validity was 75% percent, invalid time-series were calculated from fewer trials than valid time-series which might have led to increased noise and a worse estimation of oscillatory components (average number of trials: valid = 350.3, invalid = 100.6). In an additional analysis based on repeating our analysis on random samples of 25% of valid trials (1000 iterations) we found that oscillatory peaks at 3.75 Hz and 10 Hz occurred most often but were significant only in 8.6% and 9.2% of all iterations (Supplementary Figure 1).

Under the hypothetical assumption that behavior fluctuates at similar power in valid and invalid trials, this indicates that we might lack the statistical power to find significant periodic modulation in invalid RT time-series. A second potential reason for a less prominent oscillatory component in invalid trials could be the presence of additional cognitive processes related to the infrequent occurrence of the probe. Participants might be more surprised by invalid probes, leading to slower reaction times and more noise in RT time-series. Finally, it is possible that the cue led to an attentional de-prioritization of the un-cued item. This could lead either to a weaker or to a less frequent rhythmic activation of the item in memory and hence reduce its periodic impact on reaction times. Unfortunately, we cannot distinguish between these possibilities on the basis of our findings.

The analysis of the 4.6 Hz oscillatory component of valid and invalid RT time-series revealed that they fluctuated in phase opposition indicative of attention sampling both items in alternation. Theoretically both items in working memory could have been rhythmically attended independently of each other with no influence of the cue, however we consider this unlikely for the following reasons. First, valid and invalid time-series contained equal amounts of trials in which Stimulus A or Stimulus B was probed. If rhythmic sampling is initiated at stimulus onset and would remain unchanged by the cue, time-series related to Stimulus A and B should have consistent phases across all trials. Valid and invalid time-series calculated from these trials however showed opposite phases. We therefore hypothesize that attention was reset by the cue and then rhythmically sampled both items in alternation.

### Alpha rhythmic fluctuations in accuracy

Our analysis of accuracy time-series revealed a significant rhythmic modulation only in the invalid condition. Performance for un-cued probes fluctuated at a frequency of ∼9 Hz. We observed no significant anti-phasic relationship between valid and invalid accuracy time-series at 9.4 Hz or any other neighboring frequencies. Rhythmic fluctuations of in behavioral time-courses in the 10 Hz are frequently observed in behavioral studies (see VanRullen, 2016 for a review). Investigating attentional fluctuations in working memory, Pomper et. al. also found accuracy time-series to be modulated at 9.7 Hz (Pomper & Ansorge, 2021) indicating that this specific rhythmicity might be another hallmark of attentional sampling of perceived or memorized visual representations.

Most similar to our findings, a recent study on perceptual and attentional sampling found a 10 Hz oscillation in the precision parameter occurring for invalid trials exclusively (Michel et al., n.d.). The authors argued that this 10 Hz rhythm might be explained by a recently proposed model by Fiebelkorn and Kastner (2019). In this model the attentional system is hypothesized to rhythmically alternate between phases of sampling and phases of attentional shifting at a theta rhythm. During periods of attentional sampling, relevant sensory representations would be boosted. Periods of attentional shifting are hypothesized to allow the visual system to disengage from the current target of attention and become more sensitive to other relevant stimuli in the visual field. Importantly, attentional shifting is accompanied by increases in alpha oscillations that are phase locked to individual sampling/shifting theta cycles. Attentional shifts to the uncued item might therefore lead to a time-locked increase in alpha power and hence result in an alpha cyclic modulation of RT’s.

### Comparison to other studies

Our paradigm complements the findings by another study on rhythmic fluctuations in working memory in several ways (Pomper & Ansorge, 2021). Pomper & Ansorge reset internal attention via the presentation of the second working memory item whereas we used an explicit color cue. The external attentional sampling literature has found periodicities, both when using explicit cues (Fiebelkorn et al., 2013, 2018; Landau & Fries, 2012; Song et al., 2014; VanRullen et al., 2007) and when using stimulus induced attention (Chota et al., 2018; Kienitz et al., 2018; Re et al., 2019b). The difference between these two types of attentional reset however becomes more relevant in the context of working memory. Little is known about the relationship between the attentional mechanisms involved in the encoding of stimuli in working memory and the mechanisms involved in attentional prioritization after successful encoding. The fact that both types of attentional reset induce rhythmic fluctuations in behaviour however, indicates that the attentional mechanisms responsible for rhythmic sampling are involved in both encoding and subsequent prioritization of information in working memory.

Another important aspect of our study is the fact that we used multi-feature stimuli that had to be encoded as bound representations. Better accuracy and faster reaction times for cued items indicated that participants successfully used the color cue to prioritize the correct item (Figure 2AB). Specifically the effects of accuracy indicate that color-based retrieval also led to a prioritization of the orientation associated with that item. As mentioned before, the multiplexing of several multi-feature representations in working memory might serve to discretely separate features belonging to one items and features belonging to another item (Hitch et al., 2020; Nakayama & Motoyoshi, 2019; Treisman, 1996). Our study supports this claim by showing that items which are prioritized via their color feature show a facilitation of probe comparisons based on their orientation feature.

### Functional significance and neural mechanisms

There are several functional processes that could explain the rhythmicities found in our study. On the one hand, a periodic attentional refresh of working memory representations could be a fundamental requirement for maintenance. Evidence from computational modeling studies have proposed that such periodic reactivation, or “replay”, might underly working memory maintenance and could be implemented by an interplay of theta (4 to 8 Hz) and gamma (30 to 80 Hz) oscillations (Jensen & Lisman, 2005; Lisman, 1999; Lisman & Idiart, 1995). Periodic attentional refresh has also been proposed to counteract time-based decay in visual working memory. Evidence for time-based decay however has not been conclusive (Gressmann & Janczyk, 2016; van Moorselaar et al., 2015). It has been proposed that alternating activation of working memory representations might help to solve the binding problem by combining stimulus features into coherent representations (Nakayama & Motoyoshi, 2019). In our experiment this would allow the visual system to keep color features bound to the corresponding orientation features by activating the correct combination of features one at a time. This time-based multiplexing mechanism could also help to prevent merging of neural representation and might aid WM readout (Lisman & Idiart, 1995; Lundqvist et al., 2011; Miller et al., 2018,; Sandberg et al., 2003; Siegel et al., 2009).

Alternatively, visual working memory representations might only be cyclically reactivated in task contexts where they have to be compared to external visual representations e.g. during visual search or match-to-sample tasks like the one used here. This task-specific selection process might therefore serve to periodically determine which of several features or objects biases perception at any given moment without necessarily activating the representations themselves periodically. It remains an open question if representations in VWM that do not currently serve as a visual search template follow similar periodicities.

In the past years, theories have tightly linked external attentional selection with neural oscillations (Fiebelkorn et al., 2018; Fiebelkorn & Kastner, 2019; Helfrich et al., 2018; VanRullen, 2013). Even more recently the fundamental relationship between internal attention towards working memory representations and oscillatory activity has become evident (de Vries et al., 2018, 2020; Riddle et al., 2020; Sauseng et al., 2010). Current models propose that internal attention is implemented by a dynamic interplay between frontal theta oscillations, acting as a top-down control mechanism and occipital alpha oscillations that inhibit or disinhibit single representation. Our findings might be explained by a similar mechanism that modulates relative activity between two more or less equally relevant items on a shorter timescale.

### A common rhythmic sampling mechanism for external and internal visual representations

Last we would like to propose a model that could potentially explain some of the commonalities between external and internal sampling rhythms found in most recent work. Several studies have already provided convincing evidence that attention to external stimuli is fluctuating in the theta range (Fiebelkorn et al., 2013; Holcombe & Chen, 2013; Landau & Fries, 2012; Song et al., 2014). Most studies converge on a frequency of ∼4 to 6 Hz with which 2 items are sampled. Importantly it was also demonstrated that extending the number of attended locations to 3 reduces the frequency at which individual items are sampled to ∼3 Hz (Holcombe & Chen, 2013). The authors hypothesized that this observation could support the existence of a single attentional sampler in the ∼10 Hz range that has the capacity to highlight 10 items/locations per second. Our study, as well as the work from Pompers (2021) and Peters (2018) have demonstrated that this rhythmic attentional sampling process extends to information that is maintained in working memory. Importantly much evidence for the sensory recruitment hypothesis has been provided, suggesting a shared substrate between perception and working memory (Gayet et al., 2018; Rademaker et al., 2019; Scimeca et al., 2018). External and internal attention would therefore have to access representations that are encoded in highly similar areas with strongly overlapping codes. It is hence conceivable that a common attentional ‘‘master’’ sampler at 10 Hz rhythmically boost activity in specific neural representations irrespective of if they reflect information that is currently perceived or encoded in working memory. Further investigations will have to show if this model of attention holds true.

### Conclusion

In conclusion we add to the evidence that attention samples features in working memory at a theta rhythm. This rhythmic attentional facilitation alternates between two relevant items in working memory, evident in anti-phasic modulation of behavioral time-series. We hypothesize that the brain might utilize this oscillatory time-multiplexing in the face of limited attentional resources, the binding problem and/or attentional refresh required for memory maintenance. The patterns found in this study closely mimic recently found rhytmicities in external spatial and feature-based attention that were shown to be tightly linked to frontal theta and occipital alpha oscillation. Identical oscillatory mechanisms have been linked to attentional control in working memory. We hypothesize a single, theta-rhythmic attentional sampling mechanism for memorized and perceived representations.

## Supplementary Material

**Supplementary Figure 1.**
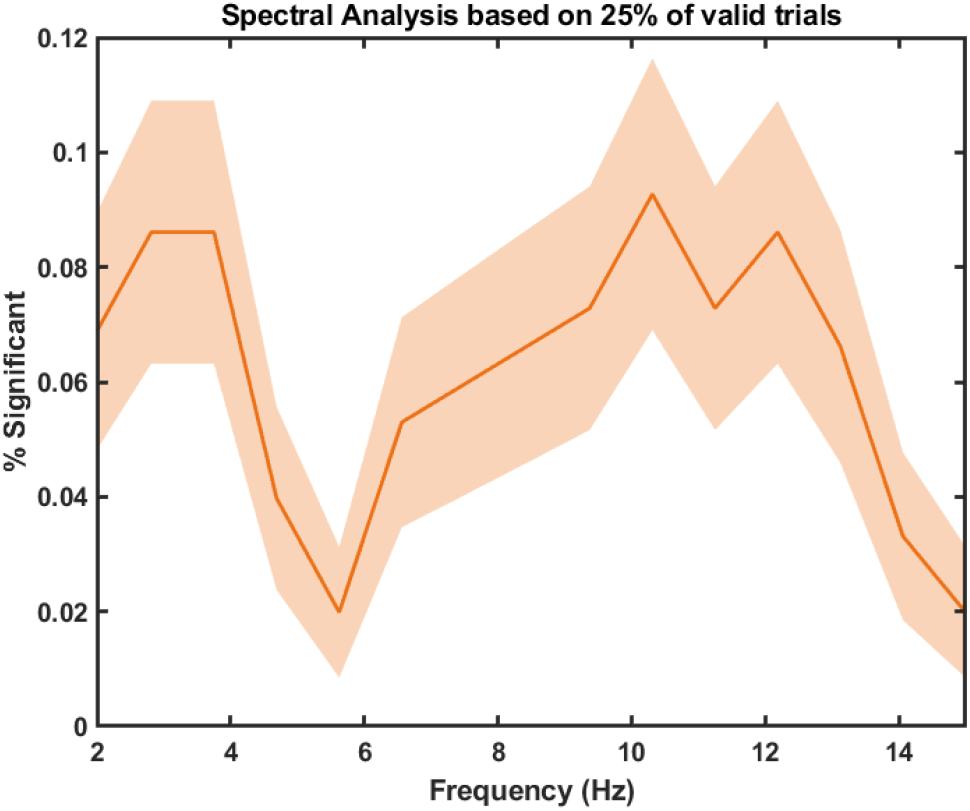
Here we tested if the lack of rhythmic fluctuations in the invalid condition were a result of a lack of statistical power due to the lower number of trials as compared to the valid condition. We reasoned that if a reanalysis of the valid condition based on the same number of trials as the invalid condition (25%) would reveal significant fluctuations, this would indicate that invalid time-series reflected weaker oscillatory activity (as opposed to a lack of statistical power). We recalculated valid time-series power-spectra 1000 times based on random samples of 25% of trials. Significant peaks in valid RT time-series in the 4 Hz range were only found in 8% of iterations, indicating that we lacked statistical power to find behavioral oscillations of equal or lower power in the invalid condition.

